# Facial masculinity is only weakly correlated with handgrip strength in young adult women

**DOI:** 10.1101/425017

**Authors:** Amanda C Hahn, Iris J Holzleitner, Anthony J Lee, Michal Kandrik, Kieran J O’Shea, Lisa M DeBruine, Benedict C Jones

**Author notes:** **Corresponding author**: Department of Psychology, Humboldt State University, 1 Harpst Street, Arcata, CA 95521. Data and analysis code are available at https://osf.io/98qf4/ and https://osf.io/chz2n/. This project was funded by ERC grants to BCJ (OCMATE) and LMD (KINSHIP).

## Abstract

**Objectives:** Ancestrally, strength is likely to have played a critical role in determining the ability to obtain and retain resources and the allocation of social status among humans. Responses to facial cues of strength are therefore thought to play an important role in human social interaction. Although many researchers have proposed that sexually dimorphic facial morphology is reliably correlated with physical strength, evidence for this hypothesis is somewhat mixed. Moreover, to date, only one study has investigated the putative relationship between facial masculinity and physical strength in women. Consequently, we tested for correlations between handgrip strength and objective measures of face-shape masculinity.

**Methods:** 531 women took part in the study. We measured each participant’s handgrip strength (dominant hand). Sexual dimorphism of face shape was objectively measured from each face photograph using two methods: discriminant analysis and vector analysis. These methods use shape components derived from principal component analyses of facial landmarks to measure the probability of the face being classified as male (discriminant analysis method) or to locate the face on a female-male continuum (vector analysis method).

**Results:** Our analyses revealed that handgrip strength is, at best, only weakly correlated with facial masculinity in women. There was a weak significant association between handgrip strength and one measure of women’s facial masculinity. The relationship between handgrip strength and our other measure of women’s facial masculinity was not significant.

**Discussion:** Together, these results do not support the hypothesis that face-shape masculinity is an important cue of physical strength, at least in women.

## Introduction

Ancestrally, strength is likely to have played a critical role in determining men’s and women’s ability to obtain and retain resources (Sell et al., 2009) and the allocation of social status (Lukaszewski et al., 2016). Being able to assess other individuals’ strength indirectly would be important to minimize the costs (e.g., injury or loss of resources) that would be incurred by engaging in competition for resources with stronger individuals. Consequently, many studies have investigated the characteristics that might function as valid cues of physical strength (Fink et al., 2007; Holzleitner & Perrett, 2016; Sell et al., 2009; Windhager et al., 2008; Van Dongen, 2014). Given the important role faces generally play in social interaction (Little et al., 2011), much of this research has investigated *facial* cues of physical strength.

Several lines of evidence suggest that human faces contain valid cues of physical strength. For example, Sell et al. (2009) found that strength ratings of face images and objective measures of upper-body strength were positively correlated in both men and women. Moreover, this pattern of results was observed in a variety of different cultures (US college students, Bolivian horticulturalists, Andean pastoralists). Relatedly, Han et al. (2017) found that dominance ratings of men’s faces were positively correlated with a composite measure of their ‘threat potential’ derived from principal component analysis of their handgrip strength, height, and weight. Using three-dimensional face images, Holzleitner and Perrett (2016) also found a weak positive correlation of upper-body strength and facial morphology in a sample of men and women.

Other studies have specifically tested whether *facial masculinity* is correlated with upper-body strength. Each of these studies used handgrip strength as their measure of upper-body strength. Fink et al. (2007) found that masculinity ratings of 32 men’s faces were positively correlated with their handgrip strength. Consistent with this result, Windhager et al. (2011) found that masculine face shape was positively correlated with handgrip strength in a sample of 26 men. By contrast with these findings, Van Dongen (2014) found that an objective measure of face-shape masculinity and handgrip strength were positively correlated in a sample of 112 women, but not in a sample of 92 men.

To date, only one study has investigated the relationship between facial masculinity and physical strength in women (Van Dongen, 2014), finding that women with more masculine faces had greater handgrip strength. The current study attempted to replicate that finding in a sample of 531 women.

## Methods

### Participants

Five hundred and thirty-one young adult women took part in the study (mean age=21.44 years, SD=3.18 years), which was part of a larger project on hormones and mating psychology (Jones et al., 2018a, 2018b, 2018c). All of the women who participated in the study were from the University of Glasgow.

### Face photography

We used a Nikon D300S digital camera with an AF Micro-Nikkor 60mm (f/2.8D) lens to take a full-face digital photograph of each woman in a small windowless room, against a constant background, and under standardized diffuse lighting conditions. Participants posed with neutral expressions. Camera settings and camera-to-head distance were held constant.

### Handgrip strength

We measured each participant’s handgrip strength from their dominant hand two times using a T. K. K. 5001 Grip A dynamometer. Following Fink et al. (2007), the highest recording from each participant (i.e., their maximal handgrip strength) was used in analyses (M=26.20 kg, SD=4.99 kg).

### Facial metrics

Sexual dimorphism of face shape was objectively measured from each face photograph using two methods: a discriminant analysis method (see Scott et al., 2010 and Lee et al., 2014 for methods) and vector analysis method (see Komori et al., 2011 and Holzleitner & Perrett, 2016 for methods). These methods use shape components derived from principal component analyses of facial landmarks to measure the probability of the face being classified as male (for the discriminant analysis method) or to locate the face on a female-male continuum (for the vector analysis method). Code for calculating discriminant and vector scores is publicly available at https://osf.io/98qf4/. An additional 50 male (Mean age=20.85 years, SD=3.01 years) and 50 female (Mean age=20.60 years, SD=1.38 years) faces (all students at University of Glasgow) were used to calculate these scores. Higher discriminant scores or higher vector scores indicate more masculine face shapes. Specific scores used in these analyses have previously been reported in Zhang et al’s (2018) study of facial correlates of women’s sexual desire and sociosexuality (the sample in the current study is smaller than Zhang et al. because handgrip strength was not measured from all women in that study).

## Results

Data and analysis code are publicly available at https://osf.io/chz2n/. Discriminant scores and vector scores were positively correlated (r=.57, N=531, p<.001). Vector scores were significantly, but weakly, positively correlated with handgrip strength (r=.09, N=531, p=.037, Figure 1). The correlation between discriminant scores and handgrip strength was not significant (r=.05, N=531, p=.30, Figure 1).

**Figure 1.**
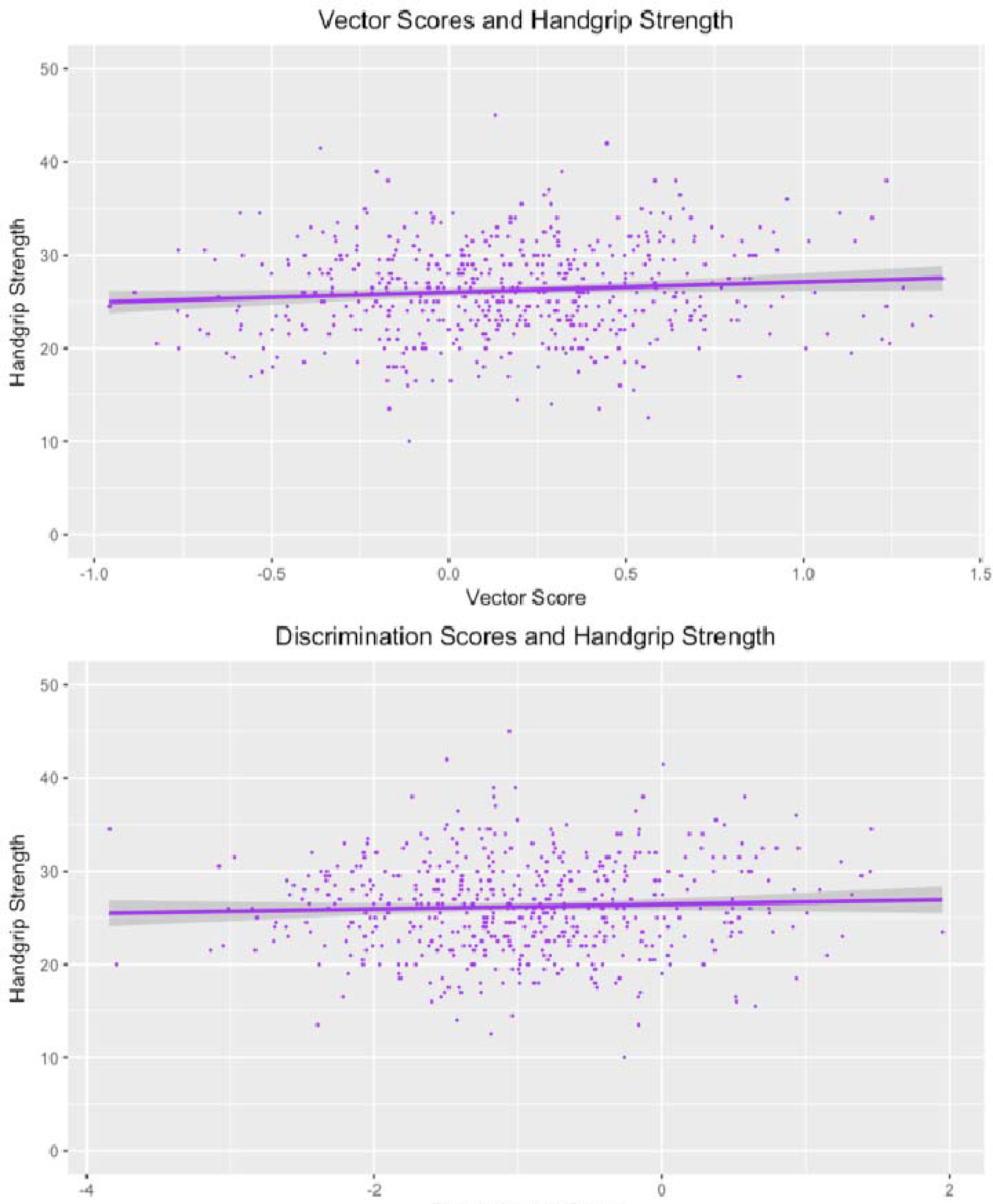
The relationships between handgrip strength (kg) and vector scores (top panel) and discriminant scores (bottom panel).

## Discussion

We tested for putative relationships between handgrip strength and two objective measures of face-shape masculinity in a sample of 531 young adult women. Although we found a significant correlation between the vector sexual dimorphism scores and handgrip strength, the correlation was very weak. In addition, we did not find a significant correlation between the discriminant scores and handgrip strength, suggesting that any potential relationship between handgrip strength and facial masculinity in our sample is not robust. The vector masculinity scores were only weakly correlated with women’s strength (r=.09) and discriminant masculinity scores were not significantly correlated with women’s strength (r=.05). These results do not support the hypothesis that morphological masculinity is an important cue for strength and strength-related perceptions of faces.

Van Dongen (2014) found that face-shape masculinity was correlated with handgrip strength in women, but not men. Our null to very small effects suggest that the correlation reported by Van Dongen (2014) for women’s face shape and handgrip strength is not robust. Both our study and Van Dongen (2014) suggest face-shape masculinity explains only a small proportion of the variance in women’s handgrip strength. Having tested only female faces, our results clearly do not speak directly to the ongoing debate of whether masculinity is a valid strength cue in men’s faces (see Fink et al., 2007; Van Dongen, 2014; Windhager et al., 2008). Given Van Dongen’s (2014) null results for handgrip strength and male faces (and the small samples in studies reporting significant correlations between these variables), we suggest that more work is needed before we can confidently conclude that there is a reliable association between handgrip strength and men’s facial masculinity.

In conclusion, despite our large sample size, we found no compelling evidence for a clear and reliable association between handgrip strength and masculine shape characteristics in women’s faces. These findings do not support the hypothesis that masculine face shapes are valid strength cues, at least in women. Our null results for strength and masculine face shapes also suggest accurate perceptions of strength from women’s faces that have been reported in previous studies are unlikely to be mediated by masculinity. Future studies using more advanced methods to capture face shape (e.g., analysis of 3D face images) could yet reveal strength-masculinity correlations even if they are not reliable in analyses of 2D images.

